# Intracellular replication dynamics of influenza A virus impose strong bottleneck effects

**DOI:** 10.1101/2025.07.18.665558

**Authors:** Ernesto Alejandro Segredo-Otero, David Gresham

## Abstract

Understanding the sources of genetic diversity in Influenza A Virus (IAV) infections is crucial for understanding the mechanisms of viral evolution and immune escape. Whereas prior studies have characterized the effects of population bottlenecks during host-to-host transmission and intrahost tissue-to-tissue dissemination, the role of intracellular replication processes on IAV genetic diversity remains largely unexplored. In this study, we used stochastic mathematical modeling to simulate the replication of genetically distinct IAV strains within individual cells and tissues. Our results reveal significant bottleneck effects within a single infection cycle of individual cells. Intracellular bottleneck effects are driven by stochastic molecular processes and lead to the expansion or elimination of neutral variants creating large-scale differences between the initial and final frequencies of genetic variants in individual cells. By expanding our findings to a population-level tissue model, we show that IAV intracellular replication reduces the effective population size, thereby diminishing the impact of selection and increasing the role of genetic drift. Our findings highlight the important contribution of intracellular replication processes to the generation of genetic diversity in IAV.

## Introduction

The genetic diversity of Influenza A Virus (IAV) plays a fundamental role in its ability to evade host immune responses, adapt to new hosts, and develop resistance to antiviral treatments. IAV evolves extremely rapidly, which allows it to persist in human and animal populations despite widespread immunity and seasonal vaccination efforts (Goldstein et al., 2015; Wang et al., 2020; Wang et al., 2020). The ability of viruses to evolve depends largely on their capacity to generate and maintain genetic diversity alongside other factors including population size and selective pressures (Brooke, 2017; Taubenberger and Kash, 2010). Understanding the mechanisms underlying the genetic diversity of IAV is essential for predicting viral evolution, guiding vaccine development, and establishing strategies to prevent the emergence of new variants (Dolan et al., 2018; Meijers et al., 2024).

As a segmented negative sense RNA virus, IAV genetic diversity arises primarily through two key mechanisms: mutation and reassortment (Shao et al., 2017). Mutations, introduced during viral RNA replication by the error-prone RNA-dependent RNA polymerase, accumulate within the viral genome and can lead to changes in viral proteins that affect antigenicity, fitness, or transmission efficiency (Muñoz-Moreno et al., 2019; Teo et al., 2025; Wang et al., 2023). Additionally, reassortment, which occurs when segments of the viral genome are exchanged between different strains during co-infection of a host cell, can produce progeny with novel genetic combinations (Ganti et al., 2022, 2021; Gong et al., 2021; Postnikova et al., 2021; Taylor et al., 2023). Together, these mechanisms generate the genetic variation observed in IAV populations and fuel the virus’s ability to rapidly adapt to selective pressures.

Genetic drift plays a crucial role in shaping the genetic diversity of viruses (McCrone and Lauring, 2018; Weaver et al., 2021). For genetic drift to play a significant role, the effective population size must remain relatively small or fluctuate markedly during specific stages of the viral life cycle—a phenomenon known as a population bottleneck. In viruses, this occurs when only a small subset of the viral population initiates new rounds of infection, resulting in random shifts in the frequencies of genetic variants (Fitzmeyer et al., 2023; Gutiérrez et al., 2012; Qu et al., 2020). In IAV, such bottlenecks can arise both during host-to-host transmission ((Lumby et al., 2020; Varble et al., 2014) and during viral spread between different tissues within an individual (Amato et al., 2022). The effects of genetic drift, especially under such bottleneck conditions, can lead to the rapid loss or fixation of genetic variants, contributing to the evolution of the viral population and influencing the outcome of infection.

Understanding the origins and consequences of viral mutations has been a central focus in IAV research. The majority of studies of mutation rates in IAV have examined polymerase fidelity (Aggarwal et al., 2010; Nobusawa and Sato, 2006; Parvin et al., 1986; Pauly et al., 2017; Suárez et al., 1992), as well as the potential trade-off between replication efficiency and fidelity (Cheung et al., 2014), and the impact of mutations on viral fitness and host adaptation (Liu et al., 2022; Muñoz-Moreno et al., 2019). In the context of population bottlenecks, research has primarily addressed their effects at the whole host level, including inter-host transmission (McCrone and Lauring, 2018; Shi et al., 2023; Sobel Leonard et al., 2017; Varble et al., 2014) and intra-host dissemination between tissues and organs (Ferreri et al., 2025). By contrast, the influence of intracellular replication dynamics on both the generation and maintenance of genetic variation remains largely unexplored.

In this study, we expanded a previously developed stochastic mathematical model of IAV intracellular replication (Heldt et al., 2015) to investigate its impact on the generation of genetic diversity. We find that strong population bottlenecks occur at the intracellular replication level and that most genome segments packaged into new virions originate from one, or very few, of the originally infecting virions, even at high multiplicities of infection (MOI). By integrating the results from the intracellular model with a population-level model, we find that the replication imbalance during early stages of infection reduces IAV effective population size, making it more prone to the effects of genetic drift, and reducing the effect of selection. The results of our model showing dramatically reduced effective population sizes is consistent with empirical observations from experiments, and demonstrates that random intracellular processes are key determinants of IAV genetic diversity.

## Results

### IAV replication imposes population bottlenecks at the intracellular level

We sought to model infection and replication in individual cells by one or more strains of IAV. Therefore, we expanded the stochastic model framework established by Heldt et al., 2015, which used a Gillespie-simulation (Gillespie, 2001) to simulate molecular events during IAV infection and intracellular replication using experimentally determined parameters. This model reproduces each molecular step of the IAV infection and replication cycle comprising (1) virion entry, (2) genome replication, (3) protein production, and (4) virion assembly and release. Within each of these steps multiple individual molecular reactions are simulated (**Figure 1**). We expanded this model to track the frequency of each genetic segment originating from different virions throughout the infection cycle. In addition to simulating the total abundance of each RNA, protein, and virion, our model records the frequency of each molecule derived from the genomes of the different infecting virions. Furthermore, it tracks the number of times each viral segment is replicated before being packaged into a virion or used in another replication reaction, enabling us to simultaneously quantify the number of new mutations generated, and the origin and history of the segment in which those mutations occur.

**Figure 1.**
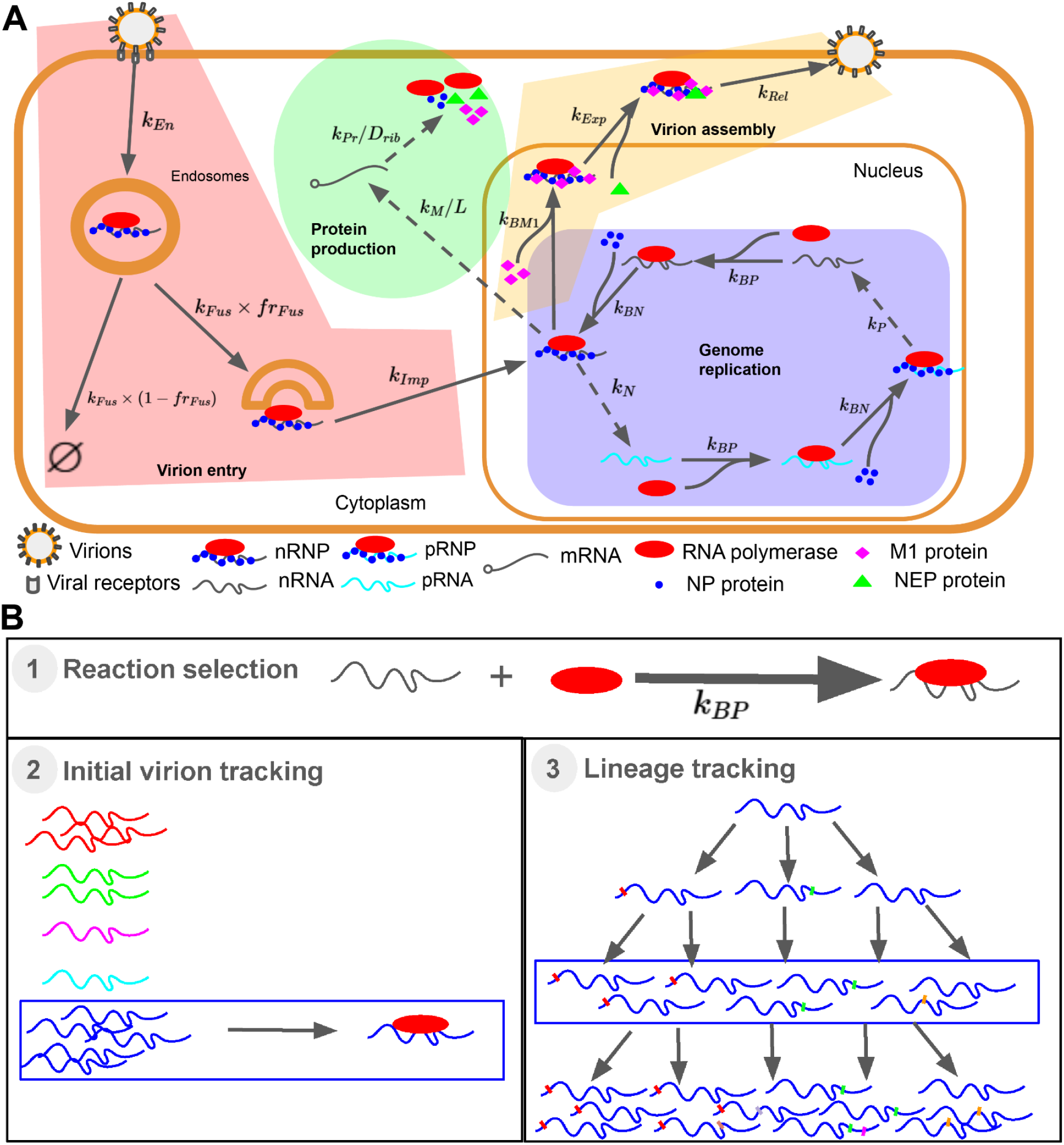
Expanded intracellular IAV replication model. **A)** Schematic representation of the IAV replication cycle, adapted from the model used by Heldt et al. (2015). All reaction rate parameters in the model are indicated: endocytosis of virions attached to receptors (*k*_*En*_); virion’s successful escape from endosomes: (*k*_*Fus*_ × *fr*_*Fus*_); virion’s unsuccessful escape from endosomes (*k*_*Fus*_ × (1 − *fr*_*Fus*_)); transport of RNPs from the cytoplasm to the nucleus (*k*_*Imp*_); replication of nRNP producing pRNA (*k*_*P*_); binding of polymerase to the naked RNA (*k*_*BP*_); binding of NP protein to the polymerase-RNA complex, forming the pRNP/nRNP (*k*_*BN*_); replication of pRNP producing nRNA (*k*_*N*_); transcription (*k*_*M*_/*L*); translation (*k*_*Pr*_/*D*_*rib*_); binding of M1 to the nRNP (*k*_*BM*1_); binding of NEP protein to the nRNP-M1 complex, resulting in export to the cytoplasm (*k*_*Exp*_); formation and export of assembled virions (*k*_*Rel*_). Parameter values and their units are presented in Table S1. **B)** Expansion of intracellular IAV replication model for studying genetic diversity. In the original model all genomes are treated as being identical. We expanded this simulation, using a three-phase strategy for each simulation step. In the first step, a standard Gillespie simulation is performed, in which both the time step (Δt) and the reaction is selected. In the second step, we randomly select the identity of the initial virion from which the genome segment originated (indicated with different colors). In the third step, we select the lineage of the genome segment, tracking how many replication rounds the genome segment has undergone.

Using this simulation framework, we first focused on the distribution of genomes originating from each initial virion throughout the infection. We found that, even in high-MOI infections, the majority of viral segments packaged into progeny virions came from only one of the infecting virions (**Figure 2A**). Even with MOI as high as 10 (i.e. infection by 10 different virions), 60%-70% of each packaged genomic segment is derived from only 1 of the infecting virions. Note, this doesn’t imply that every genomic segment is derived from the same virion. Rather, 60% of the segment 1 may be derived from virion 1, and 60% of segment 2 may be derived from virion 10, and so on.

**Figure 2.**
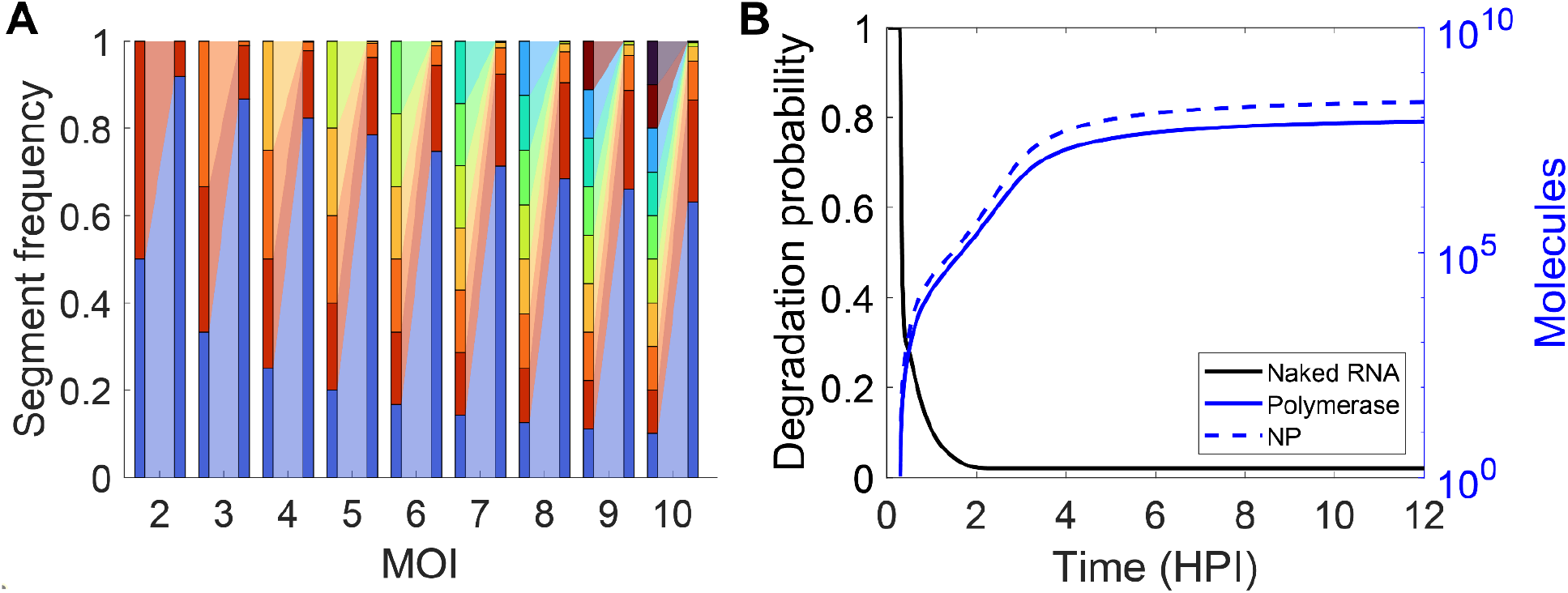
Disproportionate representation of viral genome segments following one round of IAV infection. **A)** Frequency of initial infecting (left bar) and assembled and released (right bar) genome segments originating from each unique infecting virion (indicated by color) during a single round of cell infection for MOIs ranging from 2 to 10. For every MOI the average frequency of genomic segments coming from every infecting virion, determined as 1/MOI, and the final frequency following a round of infection averaged over 1000 simulations, is shown. For example, for MOI 4, each initial segment has a frequency of 0.25. Following one round of infection, 80% of segment 1 is derived from 1 of the 4 infecting virions. **B)** Probability of RNA degradation (black) and the abundance of RNA polymerase and NP protein (blue) over time (hours post infection) for an infection with MOI of 10. Only productive infections were considered.

This dramatic skew in the fate of individual segments from infecting virions is primarily due to two phenomena that were identified by (Heldt et al., 2015) as underlying cell-to-cell variability in viral titers. First, approximately half of the infecting virions fail to reach the cell cytoplasm and are degraded in lysosomes, reducing the total number of genomes that are able to undergo replication. Second, rapid viral RNA degradation occurs within the nucleus. Each RNA-protein complex (RNP) produces naked RNAs during replication, which are highly susceptible to degradation by cellular RNAses. Once naked RNAs associate with free polymerases and NP proteins to form new RNPs, the degradation rate of RNAs is significantly reduced. Thus, the probability of degradation is calculated as the ratio of the naked RNA degradation rate to the RNA-protein binding rates, both of which depend on polymerase and NP abundance (see **Method** section for a detailed description of the model). Thus, the degradation probability evolves over the course of the infection cycle; initially this probability is high during the first hour post-infection, and it subsequently decreases as protein abundance increases (**Figure 2B**). The few RNA molecules that survive the high-degradation rate phase of the replication cycle contribute the vast majority of RNAs to the RNP population that is finally packaged into virions. Analysis of the temporal dynamics of RNA molecules shows that one or two initial virions contribute the vast majority of the total RNA molecules for each segment (**Figure S1**). The departure from the null expectation (i.e. a segment frequency of 1/MOI, which would result from every virion replicating equally), primarily due to early degradation of virions prior to entry into the nucleus and the rapid degradation of viral RNAs before the nuclear phase of the infection is established, indicates that each IAV infection event results in a strong bottleneck, amplifying the frequency of a small number of segments and eliminating the rest. Note that this imbalance in the replication of genomes from different infecting virions does not depend on the productivity of the infection, since we detect no correlation between the frequency of the dominant variant and the amount of virions produced per cell (**Figure S2**).

### IAV MOI impacts reassortment rates but not mutation rates

We sought to model the rate of reassortment and the rate of mutation. These two mechanisms are critical for the virus’s adaptability and evolution, contributing to the emergence of new strains with altered virulence, transmission capabilities, or antigenic profiles. Note, that both reassortment and mutation rate are metrics determined through assessment of the packaged and released virions from productive infections. By tracking the fate of individual viral segments and mutations throughout the intracellular replication cycle, we were able to quantify the reassortment and mutation rate under various MOIs. To estimate these rates we summarized results as the average of 1,000 simulations, which is analogous to 1,000 independent cellular infections. We find that MOI has a large impact on the rate of reassortment, with infections initiated with a higher MOI leading to a significantly increased rate of reassortment (**Figure 3A**). By contrast, the rate at which new mutations are generated is minimally impacted by MOI, decreasing slightly as the MOI increases (**Figure 3B**).

**Figure 3.**
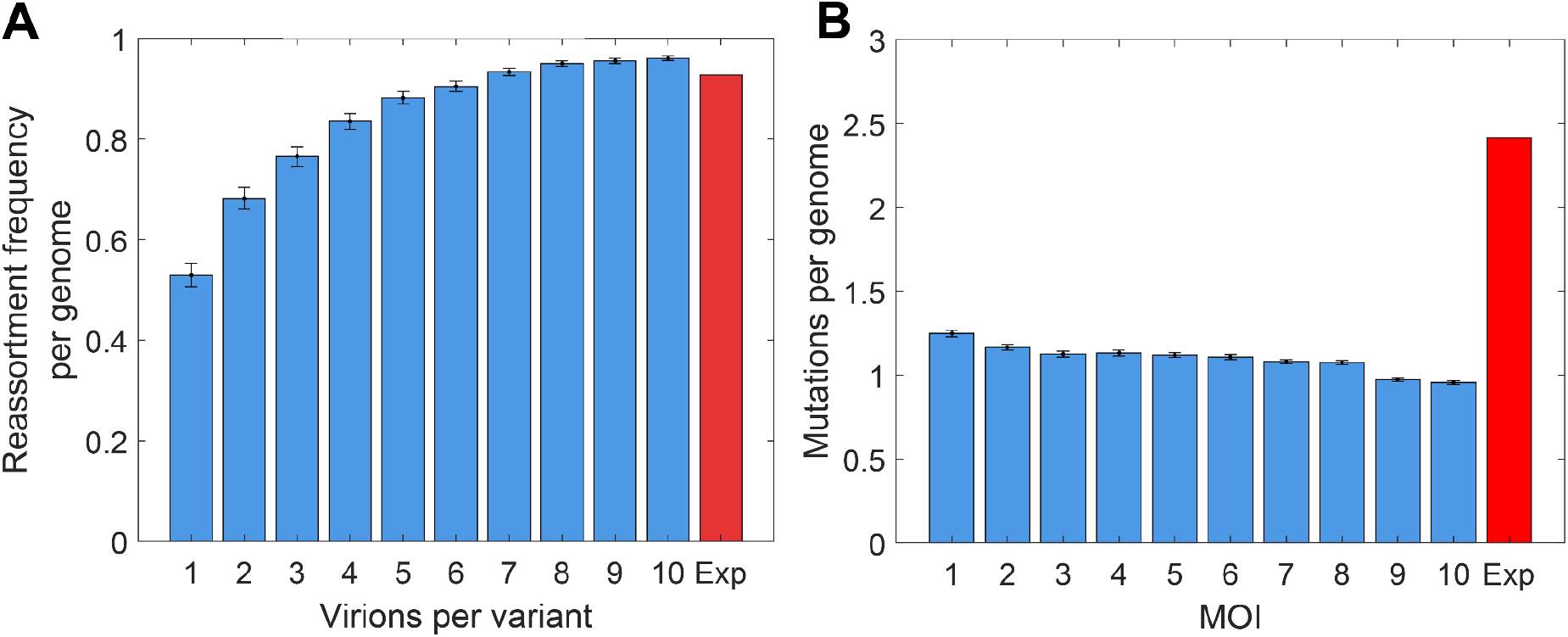
Average reassortment frequency and mutation rate as a function of MOI. **A)** Frequency of virions with reassorted genomes as a function of MOI. The red bar denotes the experimentally determined reassortment frequency between two H1N1 strains measured in (Taylor et al., 2023). **B)** Mutation rate as a function of the MOI. The experimentally determined mutation rate of H1N1 (Pauly et al., 2017) is shown in red. Bar height indicates average values and error bars indicate 95% CI. Only productive infections were considered.

To evaluate the predictive power of the model, we compared our simulated results with experimental data from the literature. Reassortment rates have recently been measured for several IAV strains, including two H1N1 strains, under controlled experimental conditions using a bulk MOI of 10 per strain (Taylor et al., 2023). A bulk MOI of 10 corresponds to an actual number of infecting virions ranging between 4 and 8 per cell (Martin et al., 2020). We found that experimentally measured reassortment rates fall within the range predicted by our model for these MOI values, aligning most closely with the simulation output corresponding to an MOI of 7 (**Figure 3A**).

To determine mutation rates using our mechanistic model we tracked the number of replication cycles each genome segment underwent prior to packaging allowing us to simulate the accumulation of mutations in each round of replication due to polymerase errors. We used the experimentally determined polymerase error rate per base of 8.3333 x 10^-6^ (Aggarwal et al., 2010; Miller et al., 2024). We modeled the probability of introducing a mutation in a segment as a Poisson process with mean α × *L* × *c*, where α is the per-base error rate, *L* is the length of the segment and *c* is the number of replication cycles. Additionally, we recorded the replication cycle history of each genome and antigenome, allowing us to estimate the probability that a given mutation becomes amplified through successive replication rounds. For instance, a mutation arising in cycle 2 is more likely to be incorporated into a larger number of progeny virions compared to one that emerges in cycle 7.

We compared simulated mutation rates for different MOIs with experimentally determined values (**Figure 3B**). IAV mutation rates have been extensively studied, although typical methods tend to be biased toward detecting mutations with minimal fitness effects. Correcting for this bias, the estimated mutation rate per genome for H1N1 is 1.8×10^-4^ mutations/site, averaging transitions and transversions (Pauly et al., 2017). Given the 13.5kb length of the IAV genome, this corresponds to an average of 2.4 mutations per replicated and packaged genome. Using our simulations we find that the mutation rate for different MOIs is between 1.24 mutations for an MOI of 1 and 0.96 mutations for an MOI of 10. The mutation rate in our model is directly related to the number of replication cycles, *c*, with increasing MOIs requiring fewer replication cycles prior to release (i.e for an MOI of 1, 7 cycles occur before release and for an MOI of 10, 5 cycles occur before release). Thus, the mutation rate decreases with higher MOI because fewer replication cycles occur prior to assembly and release. The less than two-fold discrepancy between our modeled results and experimental results is consistent with the accuracy of our method.

### Intracellular bottlenecks reduce IAV effective population size

Simulation results show that each IAV infection of a single cell functions as a genetic bottleneck, leading to stochastic amplification or reduction in the frequency of specific viral genome segments. This phenomenon, combined with the fact that some infections fail due to degradation of viral RNA or the inability of virions to escape endosomes (as previously reported by Heldt et al., 2015), suggests that the effective population size (the number of genomes or individuals from a given generation that contribute to the genetic composition of the next generation) of IAV is substantially lower than expected under a null model that assumes that all viral genomes expand equally. In this null model, the effective population size would be equal to the real (census) population size, which is simply the number of infected cells.

To expand our model to a cell population, we developed a population-level model to simulate the dynamics of viral variant frequencies across generations. We assumed a fixed number of infected cells and a constant multiplicity of infection (MOI) for each replication cycle (i.e. a single infection-replication-release cycle). We initialized the model with a set of distinct viral variants, with neutral fitness effects and low initial frequencies, mimicking the design of barcoded virus experiments (Varble et al., 2014). In each generation viral genomes were randomly sampled based on their population frequencies and assigned to cells according to the defined MOI (i.e., total number of infecting genomes equals MOI multiplied by the number of cells). In the null model, the change in frequency each generation is simply the result of binomial sampling of genomes from each cell and summing the frequency of each variant across all infected cells. The frequency of each viral variant through the generations, for the null case, is calculated as:

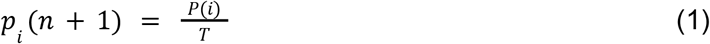

Where *p*_*i*_ is the frequency of the *ith* viral variant, *T* is the total number of infections, and *P*(*i*) is a random number sampled from a multinomial distribution with mean *p*_*i*_ (*n*) × *T*, such that 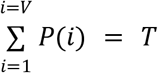 for *V* total variants. In our model we calculate *P*(*i*) by multiplying each number by their relative abundance intracellularly as a result of the infection, and the amount of virions produced by that cell. If the infection is not successful, we set the contribution of that cell to 0.

Using this approach, we compared the effective population size under the null scenario—in which all cells are effectively infected, and all genomes replicate equally—with our simulation results. To calculate the effective population size, the expected change in the frequency of a variant allele (p_i_) due to genetic drift is given by the formula (Charlesworth, 2009):

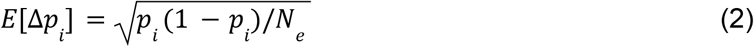

where *p*_*i*_ represents the frequency of the *ith* allele, and *N*_*e*_ is the effective population size. Using this relationship, we computed *N*_*e*_ using either the allele frequency changes observed in our simulations or those predicted by the null model, which allowed us to quantify how stochastic events during the IAV replication cycle impact the effective population size relative to theoretical expectations. We observe that *N*_*e*_ in the null case is simply related to the census population size (i.e the total number of infections), whereas the IAV model results in an *N*_*e*_ that is between 1-2 orders of magnitude lower than the census size (**Figure 4A**).

**Figure 4.**
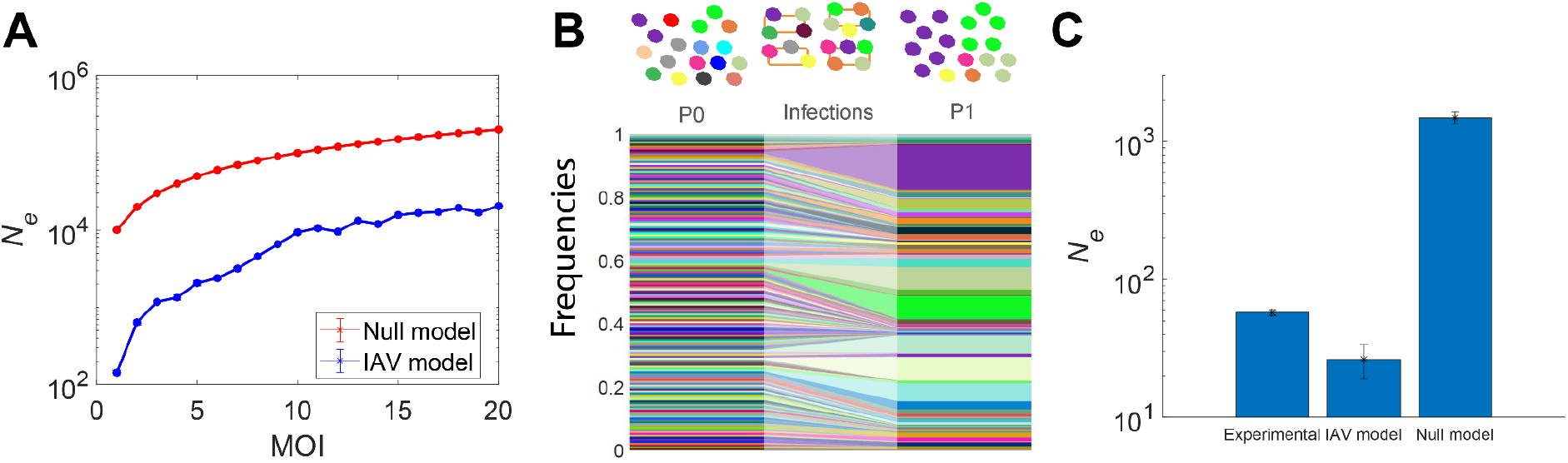
IAV intracellular bottlenecks affect the effective population size of IAV. **A)** Comparison of *N*_*e*_ between the null model and the IAV intracellular model following a single generation. **B)**. Simulation of barcoded viral experiment. We initiated infections with the same number of variants as in their experiment, all equally distributed, and used their frequency before and after IAV infection of the whole cell population, used to estimate *N*_*e*_. **C)** Estimation of the effective population size of an IAV plate infection of MDCK cells using equation 1, from the experimental data presented in (Varble et al., 2014), and our estimation using the IAV model or the null model. For both models, we estimated the number of infected cells per generation based on the viral titer, the number of virions per cell predicted by the model, and the IAV virion degradation rate estimated in (Baccam et al., 2006) (see **Methods section**).

We tested whether the discrepancy between the predictions of the null model and our IAV population size model explain experimental observations of IAV genetic diversity. Varble et al., (2014) employed barcoded IAV infections in cell culture to track the frequency of neutral variants before and after infection. In addition, they measured viral titers at 12, 24, 36, and 48 hours post-infection (hpi). We simulated this experimental procedure using our population model, with a set of initial variants equal to the total number of barcoded variants in their experiment (**Figure 2B**). Using the viral titer measurements, we estimated the number of infected cells per generation by analyzing the changes in viral load between time points—used here as proxies for viral generations—while adjusting for virion degradation rates and the fraction of unsuccessful infections predicted by our intracellular model using an MOI of 1 (as experimentally determined by plaque assay; see **Methods** for details). For the first generation, we assumed an MOI of 1, given that the population-level MOI was 0.01, implying that most infections were initiated by a single virion. For subsequent generations, we assumed a high local MOI (i.e. an MOI of 20), reflecting the focal nature of viral spread, which leads to elevated local concentrations of virions around susceptible cells. Incorporating barcode frequency data and our estimates of cell infection rates and MOI, our model predicts an effective population size of 25 ± 6, compared to the experimental value of 57 ± 3. By contrast, the null model dramatically overestimates the effective population size, yielding 1.18 × 10^3^ ± 137.14 (**Figure 4C**). The approximately two-fold difference between our estimate of *N*_*e*_ and that derived from experimental data again points to the accuracy of the model.

### Intracellular bottlenecks reduce the effectiveness of selection

To assess the impact of intracellular bottlenecks on the efficiency of directional selection, we extended our population-level model to simulate the dynamics of advantageous mutations. Specifically, we tracked the frequency of a single beneficial variant across multiple generations under varying MOI and population sizes (i.e., number of infected cells). To incorporate selection, we modified our population model to account for the fitness advantage of a beneficial variant by quantifying the frequency of each variant using :

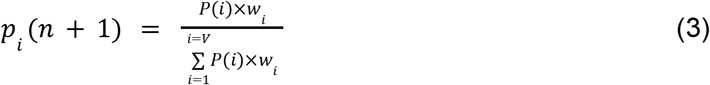

Where *w*_*i*_ is the relative fitness of each viral variant. To evaluate the impact of selection for different MOIs we simulated a total number of 10^5^ virions per generation. This was achieved by adjusting the number of infected cells accordingly for each MOI, ensuring that differences in outcomes would reflect the effect of intracellular dynamics and not changes in population size. We introduced a high-fitness variant (HFV) with fitness effects varying between 1.5 and 5 (i.e. a 50% - 500% growth advantage per generation), whereas all other variants had neutral fitness effects. We determined the frequency of the HFV after 28 generations—equivalent to 14 days of infection assuming a 12-hour replication cycle—and recorded the proportion of simulations in which the variant reached fixation (defined as a frequency *≥* 0.999). Using our model we find that the probability of fixation is low when the MOI is 1 and fitness advantage is modest (**Figure 5A**). Increasing either MOI or fitness effect results in an increased probability of fixation (**Figure 5B**). However, even at the most extreme values of MOI and fitness effect, the probability of fixation never exceeds 0.9.

**Figure 5.**
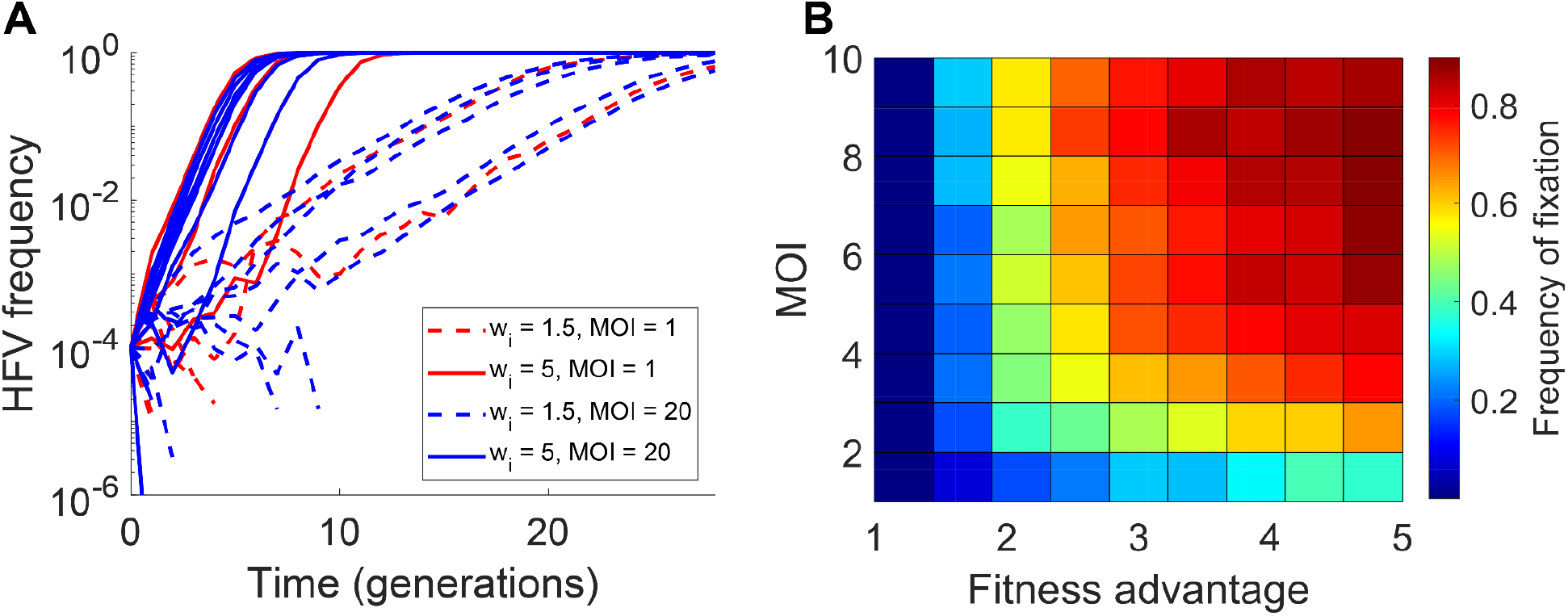
Effect of IAV bottlenecks on positive selection. **A)** Frequency of the high fitness variant (HFV) over generations. Data are shown for 10 different simulations for each parameter combination. **B)** Fraction of times that the high-fitness variant reaches a frequency higher than 0.999 in a 14 days infection cycle. In simulations, we used an initial frequency of 10^-4^ and a total of 10^4^ virions per generation.Data are the average of 1,000 simulations.

## Discussion

In this study, we used stochastic simulations to study how intracellular replication of IAV impacts several evolutionary properties features including effective population size, directional selection, and mutation and reassortment rates. Building on an experimentally-informed stochastic model of IAV infection (Heldt et al., 2015), we expanded the model to track the replication of each individual genome derived from individual infecting virions and the replication history of each genomic segment. We found that each IAV cellular infection behaves as a bottleneck event randomly amplifying or reducing the frequency of the infecting variants. This phenomenon is mainly explained by two factors: 1) the degradation of virions in endosomes and their failure to reach the nucleus and 2) the stochastic degradation of naked viral RNAs in the nucleus. These two factors also the primary determinants of the cell-to-cell heterogeneity in viral titiers (Heldt et al., 2015). Our model reproduced experimental observations including the H1N1 reassortment rate, and the observed IAV mutation rate. We demonstrated that intracellular bottlenecks drastically reduce the effective population size, especially at low MOI, thereby weakening the effects of positive selection.

Our findings highlight the contribution of cellular RNA degradation processes to IAV genetic diversity. Free unadenylated RNAs in the nucleus of mammalian cells are degraded by the exosome, with the nuclear exosome targeting (NEXT) complex playing a central role in directing these RNAs to this degradation pathway (Gerlach et al., 2022; Lubas et al., 2011). Although the exosome’s main function is to eliminate spurious host transcripts, it also acts as an effective defense against RNA viruses, making RNA stabilization crucial for viral survival. This is achieved by IAV mainly through interactions with NP proteins (Lee et al., 2017; Turrell et al., 2013). In the earliest stages of an infection, the relative paucity of NP proteins means that viral RNAs are highly likely to be degraded thereby altering the population of viral RNAs that ultimately contribute to the released virions and thereby genetic variation.

Electron microscopy studies show that up to 80% of IAV virions carry the complete set of eight genomic segments (Nakatsu et al., 2016), a finding seemingly at odds with experimental data suggesting that single-virion infections rarely replicate all eight segments (Brooke et al., 2013; Jacobs et al., 2019). However, the original model by Heldt et al., (2015) reconciled these observations, showing that although infecting virions may carry all segments, stochastic degradation of RNAs often prevents full replication of all 8 segments within a cell leading to unsuccessful infections. Our work shows that this random degradation not only contributes to failed infections (i.e. no virion release) at low MOI but also leads to imbalanced amplification of segments from different virions at high MOI, resulting in the amplification or loss of variants. Consequently, each IAV infection, even when successful, acts as a bottleneck reducing the number of genomes that ultimately contribute to the viral progeny.

Genomic reassortment is a key driver of IAV evolution and has been responsible for the emergence of pandemic strains (Garten et al., 2009). Reassortment can only occur when at least two distinct virions from different strains coinfect not only the same host but also the same cell. Recent findings have shown that reassortment in IAV is strain-dependent (Taylor et al., 2023); however, this dependency is not explained by genetic similarity between strains, but rather by inherent differences in reassortment capacity. Investigating the replication dynamics of IAV may help identify the factors underlying this strain-specific behavior. Our model accurately reproduces the experimentally observed reassortment rate of H1N1 IAV and suggests that this rate is primarily influenced by the fraction of virions that escape the endosomes and the ability of their RNAs to evade degradation and replicate. Quantifying these parameters for different IAV strains may be informative for predicting reassortment potential.

Although the proximate source of genetic mutation in IAV is polymerase fidelity, the entire replication process contributes to the observed variation in virions. In RNA viruses, two replication strategies are possible: stamping machine and geometric growth. The stamping machine model operates when the replication rates of genomes and anti-genomes are highly imbalanced, producing very few anti-genomes that serve as templates for most genome synthesis. By contrast, geometric growth involves equal replication rates for genomes and anti-genomes, leading to multiple rounds of replication for both strands (Martínez et al., 2011; Sardanyés et al., 2012; Schulte et al., 2015) and is associated with a higher accumulation of mutations (Sardanyés et al., 2009; Schulte et al., 2015). In the case of IAV, the replication rate of nRNA (genomic segments) is approximately ten times higher than that of pRNA (anti-genomes), placing it in an intermediate position between these two models. The IAV model shows that the experimentally observed mutation rate (Pauly et al., 2017) can be explained by the combined effect of error prone replication by RNA polymerase and intracellular replication dynamics.

Population bottlenecks occur at multiple stages of the viral life cycle, particularly in multicellular hosts (Amato et al., 2022; Ferreri et al., 2025; Lumby et al., 2020; Martin et al., 2024; McCrone and Lauring, 2018; Shi et al., 2024; Sobel Leonard et al., 2017; Varble et al., 2014; Zwart and Elena, 2015). These transmission events reduce the effective population size, and variation in genetic composition are often used to estimate the number of virions that successfully establish infection and act as founders of the new population (Ferreri et al., 2025; Shi et al., 2024; Sobel Leonard et al., 2017). In IAV, inter-host transmission bottlenecks have been estimated to involve as few as one or two virions per new host (McCrone and Lauring, 2018). Our results show that the IAV intracellular replication cycle also acts as a bottleneck reducing the effective population size by at least a factor of ten and thereby impacting the genetic diversity and evolution of IAV.

## Methods

### IAV intracellular replication stochastic model

Our stochastic model is based on the framework developed by (Heldt et al., 2015), which used a Gillespie simulation approach to simulate the intracellular replication of IAV, using experimentally determined parameters. The original model does not distinguish between different virions or genomes during replication, effectively considering all viral genomes as identical in terms of genetic composition. To study genetic diversity we sought to develop a model capable of tracking the origins of genomes packed into virions, identifying which of the infecting virions they came from. Treating each viral molecule as an independent species in the Gillespie simulation would require expanding the stoichiometric matrix (the matrix that relates the subtracts and products of every reaction) by a factor of MOI^2^, as each species and reaction would need to account for all possible interactions. Furthermore, as we are also interested in tracking how many replication cycles each genome undergoes before being packaged, we differentiate between viral genomes and antigenomes based on their replication cycle, setting maximum cycle number *MaxC = 30* (though no genome exceeded 12 cycles in our simulations). This would result in an unmanageably large final matrix of size *S*_*T*_×*MOI*^2^×*MaxC*^2^ ×*R*_*T*_ ×*MOI*^2^×*MaxC*^2^, in which *S*_*T*_ and *R*_*T*_ are the total number of species and reactions in an MOI 1 infection, respectively. To mitigate this, we distributed information about the total number of molecules per species, per initial virion, and per replication cycle across three distinct matrices. We retained the standard state vector, which records the number of molecules per species (*S*) at each time step. Additionally, we introduced two matrices: *S*_*M*_, with *S*_*T*_ rows per MOI columns, recording the number of molecules originating from each initial virion (which we refer to as MOI index) and their products, and *S*_*C*_, with *S*_*T*_ × *MOI* rows and *MaxC* columns, which track the number of molecules originating from each initial virion or their products at each specific replication cycle. Every reactant produces molecules corresponding to its same MOI index and replication cycle, with the exception of genomes produced by antigenomes, which generate genomes corresponding to their cycle plus 1. We then adapted the Gillespie method to simulate the different reactions involving molecules from different virions and replication cycles in a three-step process (**Figure 1B**).

First, a standard Gillespie simulation step is performed, using the vector of number of molecules per species (*S*) and the standard stoichiometric matrix (*M*). We calculate the propensities of the reactions using the vector *S*, a reaction matrix (*M*_*R*_), which consists of a matrix of the size of *M*, containing the reactants of every reaction, instead of the stoichiometric balance, and a parameter matrix (*M*_*P*_), which consists of a matrix of *P* (number of different parameters) rows per *R*_*T*_ columns. The propensities vector (*a*) is calculated as:

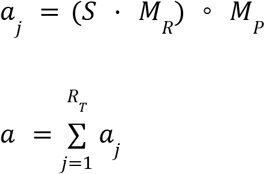

The time step and the reaction occurring is determined using the standard Gillepsie method, consisting of using two uniformly distributed random numbers, that are determined by:

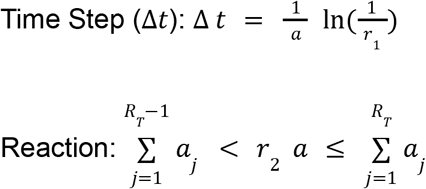

And with the specific *jith* that satisfied that condition, we updated the vector *S*.

Second, we calculate the propensities of the reaction *j* for every species of the matrix *S*_*M*_, and use another ‘*r*’ random number to decide which of the molecules from the different MOI indexes is the performing the reaction.

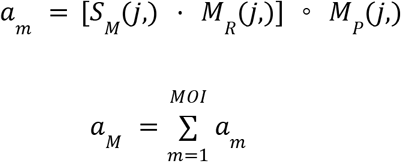

Where *S*_*M*_ (*j*,), *M* _*R*_(*j*,) and *M*_*P*_ (*j*,) are the reactant species, stoichiometric indexes, and parameters of reaction *j* respectively. The MOI index is then determined as:

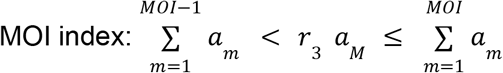

And so, we updated the matrix *S*_*M*_ with the *mth* MOI index that satisfied the equation.

Third, we use *S*_*C*_ to calculate the propensities of the reaction for every different cycle of reaction *j* and MOI index *m*, and proceed identically to determine which molecule performs the reaction.

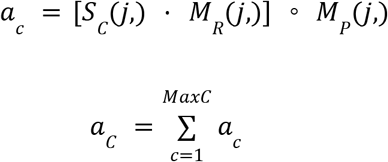

Replication cycle: 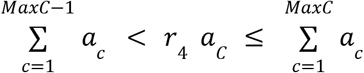

With which we updated the matrix *S*_*C*_ with the *cth* replication cycle that satisfied the equation.

For each virion produced, we perform steps 2 and 3 for each of the 8 genomic segments, with the probability of each segment being packaged at a given MOI index and replication cycle being proportional to the concentration of RNP in the cytoplasm. Importantly, we do not introduce any bias that would preferentially increase the likelihood of packaging segments from the same MOI index. Instead, the probability of packaging each segment is determined solely by its concentration.

### Modeling complex reactions

Some reactions do not follow a simple mass action behavior requiring more detailed modeling. In line with (Heldt et al., 2015), we assumed that the translation of the IAV RNA-dependent RNA polymerase (*P*_*Rprd*_) depends on the mRNA of the first three genomic segments (PB1, PB2, and PA). Specifically, the translation rate of *P*_*Rprd*_ is governed by the concentration of the least abundant mRNA among these segments. The translation rates for proteins M1, M2, and NEP are determined by the abundance of the mRNA from segments 7 (M1 and M2) and 8, scaled by the splicing factors 1 − *F*_*Spl*7_, *F*_*Spl*7_ and *F*_*Spl*8_ respectively. Additionally, each translation rate is divided by a factor, *Drib*, which accounts for the distance between adjacent ribosomes on the same mRNA strand.

Following the model of (Heldt et al., 2015), we assumed that virion release is dependent on the cytoplasmic concentration of viral ribonucleoproteins (*RNP*_*C*_) and asymptotically dependent on the protein concentration, following a Michaelis-Menten function as:

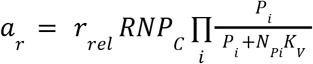

Where *r*_*rel*_ is the virion release rate, *RNP*_*C*_ is the concentration of the least abundant RNP in the cytoplasm, *P*_*i*_ is the protein concentration, *K*_*V*_ is the protein concentration at which the virion release is half maximal, and *N*_*Pi*_ is the number of proteins of each type per virion. Finally, we also incorporated the regulation of viral transcription described in (Martínez-Alonso et al., 2016; Rodriguez et al., 2007, p.; Rüdiger et al., 2019), (Rodriguez et al., 2007) and (Martínez-Alonso et al., 2016), that assumes that the free RNA-dependent RNA polymerase *P*_*Rprd*_ degrades the cellular RNA polymerase II, reducing the mRNA transcription rate, leading to the expression:

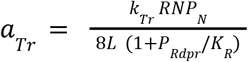

Where *k*_*Tr*_ is the transcription rate, *L* is the length of the mRNAs and *K*_*R*_ is for the concentration of polymerase (*P*_*Rdpr*_) at which the transcription rate is half inhibited.

### Computational performance approximations

We implemented a series of computational strategies to enable a large number of simulations to be performed in a short time. First, we incorporated the tau-leaping method, a well-established approach for accelerating Gillespie simulations ((Gillespie, 2001);(Cao et al., 2007)). We employed the step size selection algorithm proposed by (Cao et al., 2007), which distinguishes between critical and non-critical reactions. Critical reactions are those that cannot be triggered more than a specified number of times without generating negative amounts of molecules. Following their guidelines, we set this threshold at 10. For critical reactions, the time step (Δ*t*_*C*_) and reaction determination are handled as in the classic Gillespie simulation, while non-critical reactions are simulated by triggering them at a specific non-critical time step ‘τ_*NC*_ ‘,, which is determined based on the method outlined by (Cao et al., 2007). Once both time steps (Δ*t*_*C*_ and τ_*NC*_), are determined, the script operates as follows:

1. If τ_*NC*_ < Δ*t*_*C*_
  a. Δ*t* = τ_*NC*_
  b. No critical reaction occurs during this time step.
  c. A random number of triggers occur for each of the non-critical reactions, sampled from a poisson distribution with mean (τ_*NC*_ × *a*_NC_), being *a*_NC_ the propensities of the non-critical reactions.
  d. For every reaction that is triggered, we distribute that many numbers of reactions among all of the MOI indexes and replication cycle as defined above for steps 3 and 4 of our simulation method.
2. If τ_*NC*_ > Δ*t*_*C*_
  a. Δ*t* = Δ*t*_*C*_
  b. A standard Gillespie step is performed using only the subset of critical reactions.
  c. A random number of triggers occur for each of the non-critical reactions, sampled from a poisson distribution with mean (Δ*t*_*C*_ × *a*_NC_), being *a*_NC_ the propensities of the non-critical reactions.
  d. For every reaction that is triggered, including the critical reaction, we distribute that many number of reactions among all of the MOI indexes and replication cycle as defined above.

To avoid unnecessary applications of the approximation, we only perform tau-leaping when the new time step (i.e., the smallest value between Δ*t*_*C*_ and τ_*NC*_) is at least 10 times larger than that of the standard Gillespie simulation. Otherwise, we proceed with a normal Gillespie step. If, during the tau-leaping reactions for the MOI and replication cycle distribution, a negative value arises due to the distribution of a high number of reactions, we set the corresponding MOI index or replication cycle to 0. The number of negative molecules is then redistributed among the remaining indexes or cycles, and the propensities are recalculated with the updated zero value.

We also introduced mathematical approximations to handle species that act as fast chemical switchers in the simulations—that is, species that change states much faster than the other reactions. In our model, this phenomenon potentially occurs at three different steps in the replication cycle, involving three species that are created and consumed rapidly. During genomic replication, RNPs generate naked RNA including genomic and antigenomic RNA. These RNA species are highly unstable, as naked RNA is rapidly degraded in the cell. To protect this RNA, it binds it to polymerase and N proteins forming (RNPs). As a result, each naked RNA will either be quickly degraded or bound to a polymerase. These RNA-Polymerase (RNA-Pol) complexes are also subject to degradation or will bind to a certain number of N proteins (depending on RNA length) to form RNPs. Because these processes depend on protein concentrations, which can reach extremely high levels, their reaction rates are substantially higher than the replication rate (i.e., the generation of naked RNA by the RNPs). To avoid simulating an excessive number of small time steps and given that the only possible fates for naked RNA are either RNP formation or degradation, we assumed that the replication reaction is the limiting step in RNP formation. We then calculated the propensity for RNP formation by using the propensity of naked RNA generation, divided by the probability of the naked RNA surviving and forming an RNA-Pol complex without being degraded in the process. For each replication reaction triggered, the probability that the generated naked RNA survives and forms an RNA-Pol complex is determined by the ratio of the binding reaction rate to the sum of the binding and degradation reaction rates:

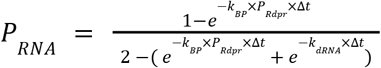

Where *k*_*B*P_ and k _*dRNA*_ are the rate of binding of the polymerase and the degradation rate, respectively. In a similar way, the probability of the complex RNA-Pol becoming an RNP is calculated as:

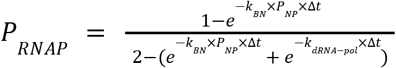

Where *k*_*B*N_ and k_d*RNA*−*po*l_ are the rate of binding of the polymerase and degradation rate, respectively, and *P*_*NP*_ is the N protein concentration. This way, we eliminated from our simulations the species of naked RNAs and RNA-Pol complex, and set the propensities of the RNP production as the ones of the naked RNA replication, multiplied by both probabilities. Each time a replication reaction is triggered, it not only produces an RNP directly, but also consumes one polymerase and as many NPs as required for the corresponding segment. We applied the same approximation to the export of RNPs from the nucleus to the cytoplasm, a crucial step for genomes to be packaged into virions. RNPs are first bound to M1 proteins and then to the NEP protein for export to the cytoplasm. Since the RNP-M1 complex in the nucleus is an intermediate state whose only fate is either degradation or export, we eliminated this intermediate state and applied the same approach as for naked RNA and RNA-Pol complexes. The propensity for export is then calculated as the propensity for M1 binding, multiplied by the probability that the RNP-M1 complex will be exported without being degraded. This probability is calculated as:

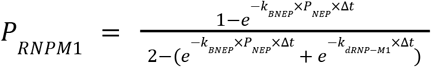

Where *k*_*B*N*E*P_ and *k*_*dRNA*−*M*1_ are the rate of binding of the polymerase and the degradation rate respectively and *P*_*NEw*P_ is the NEP protein concentration.

Finally, we observed that, in a large number of simulations, the limiting step was often the M1 binding process, as it consumes an RNP and is susceptible to being a critical reaction dependent on protein concentration. To reduce computation times, we implemented a quasi-steady-state approach for the concentration of nRNPs. We calculated their concentration as the ratio of the propensities of the reactions that form them (replication) and consume them (degradation and M1 binding), and used a deterministic approach to model the formation of exported nRNPs to the nucleus. This concentration was then distributed across MOI indices and replication cycles in proportion to the distribution from the previous step. We set a threshold for triggering this approximation, typically 0.001 min, such that:

- If Δ*t* < time limit
  - 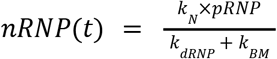
  - *nRNPM1*(t) = *nRNPM1*(t−1) + *k*_*BM*_ × *nRNP*(*t* − 1) × Δ*t*
  - a_m_ (nRnPs) = 0

The last step sets the propensities of all reactions involving nRNPs to zero, and proceeds to calculate the new time steps of the Gillespie and tau-leaping methods. If those new time steps are still lower than the threshold, we discard the approximation.

### Population level model

We developed a population-level model that incorporates the imbalance in the amplification of genomic segments during the intracellular replication of IAV. We used a Wright-Fisher-like model with a constant number of infected cells and MOI per generation, where each infected cell either increases or decreases the proportion of each infecting viral genome based on the results from the intracellular model. In the absence of the stochastic steps of IAV replication, we assumed that every infecting virion contributed equally to the outcome of the infection and that each cell produced an equal amount of virions. We referred to this scenario as the “null case”, and the evolution of the frequency of each viral variant under these conditions follows the equation:

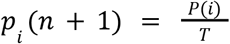

Where *p*_*i*_ stands for the frequency of the *ith* variant, *T* for the total number of genomes per generation and *P*(i) is the number of times each variant is selected to infect a cell, sampled from a multinomial distribution with mean *p*_*i*_ (*n*) × T, set so that 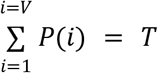, with *V* total number of variants. In the case of IAV, the contribution of each of the *P*(i) infections that every viral variant performs are weighted by their intracellular frequency after the infection occurs, and the amount of virions that cell produces. Unsuccessful infections contributed with 0. For the sake of computational efficiency, we did not perform a simulation of the viral infection during the population level simulations, but instead we pre-generated a set of 1000 simulations per MOI, and randomly assigned one of each of these results to each infected cell.

Natural selection was incorporated into the population model by weighting the contribution of each variant by its relative fitness value. In the null case, the change in variant frequency across generations then follows the updated equation:

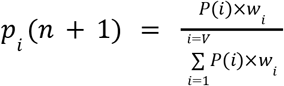

Where *w*_i_ represents the relative fitness of each viral variant. As in previous approaches, we modeled changes in variant frequency during IAV infection based on intracellular performance. Importantly, we did not alter the intracellular behavior of any variant according to its fitness; instead, all variants were treated as neutral within the intracellular model. Thus, the fitness advantage simulated in our framework stems exclusively from extracellular factors—specifically, enhanced infectivity and/or virion stability. Such advantages could arise, for example, from improved immune evasion mechanisms of the virions, which may result, for example, from a greater ability to evade the immune system.

### Modeling barcode tracking experiment

As our model is specifically parameterized for H1N1 infection in MDCK cells, we restricted our analysis to the corresponding data from this cell line from (Varble et al., 2014). To enable an accurate comparison, we needed a precise estimate of the total number of infected cells per replication cycle (approximately every 12 hours). One of the outputs of our model is the average number of virions produced per infected cell, which allowed us to estimate the number of infected cells at each time point by dividing the measured viral titers by the model-predicted virions per cell. Since the titers in (Varble et al., 2014) were obtained via plaque assays—which are performed under low MOI—we adjusted the titers by multiplying them by the inverse of the fraction of unsuccessful infections predicted by our model at MOI = 1. Additionally, we multiplied the titers by 2 to account for the 2 mL volume typically used in 6-well plates (as the titers are reported in PFU/mL). Lastly, we corrected for virion degradation using the reported IAV half-life of 3 hours from (Baccam et al., 2006). Taking all these factors into account, the estimated number of infected cells per generation is given by:

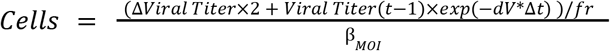

Where Δ*Vira*l Titer is difference between the viral titer from a period of 12 hours, *Vira*l Titer(*t* − 1) is the viral titer of t - 12 hours, *dV* = − *log*(0. 5)/(*v*i*rions* h*a*l*f* − *lif*e), *f*r is the fraction of successful infection of MOI 1 infections, and β is the average number of virions produced by cells of the corresponding MOI. As the infections were produced at an initial MOI of 0.01, but IAV produces focal infections that increase locally the MOI, we assumed MOI = 1 for the first infection cycle (first 12 hours), and MOI = 20 for the rest. With the viral titer data of (Varble et al., 2014), and these approaches, we calculated the total number of infected cells in every generation, which sums to 1,329,918, a value the is consistent with the 1.2 ×10^6^ cells that are expected to be in the 6-well plates used for the infections in their experiments.

### Computer code availability

All code for performing simulations was written in Matlab and is available on Github (https://github.com/GreshamLab/IAV_intracellular_model). Simulated data and the barcoded infections data from (Varble et al., 2014), used in the population level model, are uploaded in https://osf.io/f9e6v/.

## Acknowledgements

We thank members of the Gresham lab for valuable discussion and comments on the manuscript. Funding for this research was provided by NIGMS (R35GM153419), NIAID (R01AI140766 and R01AI170112), and the US - Israel Binational Science Foundation (2021276).

## Supplementary information

**Figure S1.**
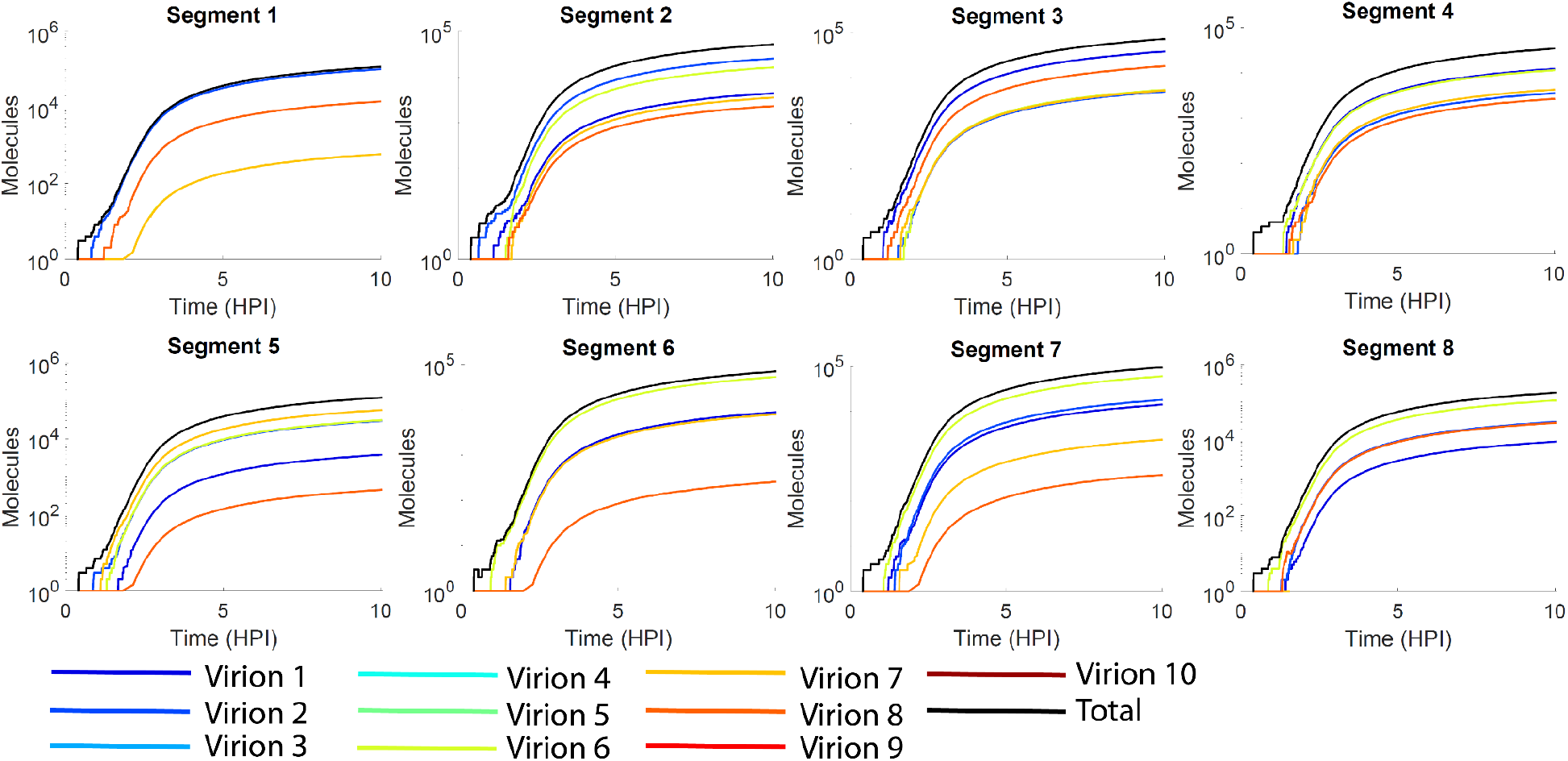
Dynamics of increase in number of molecules for each genome segment from each infecting virion for an infection with MOI of 10.

**Figure S2.**
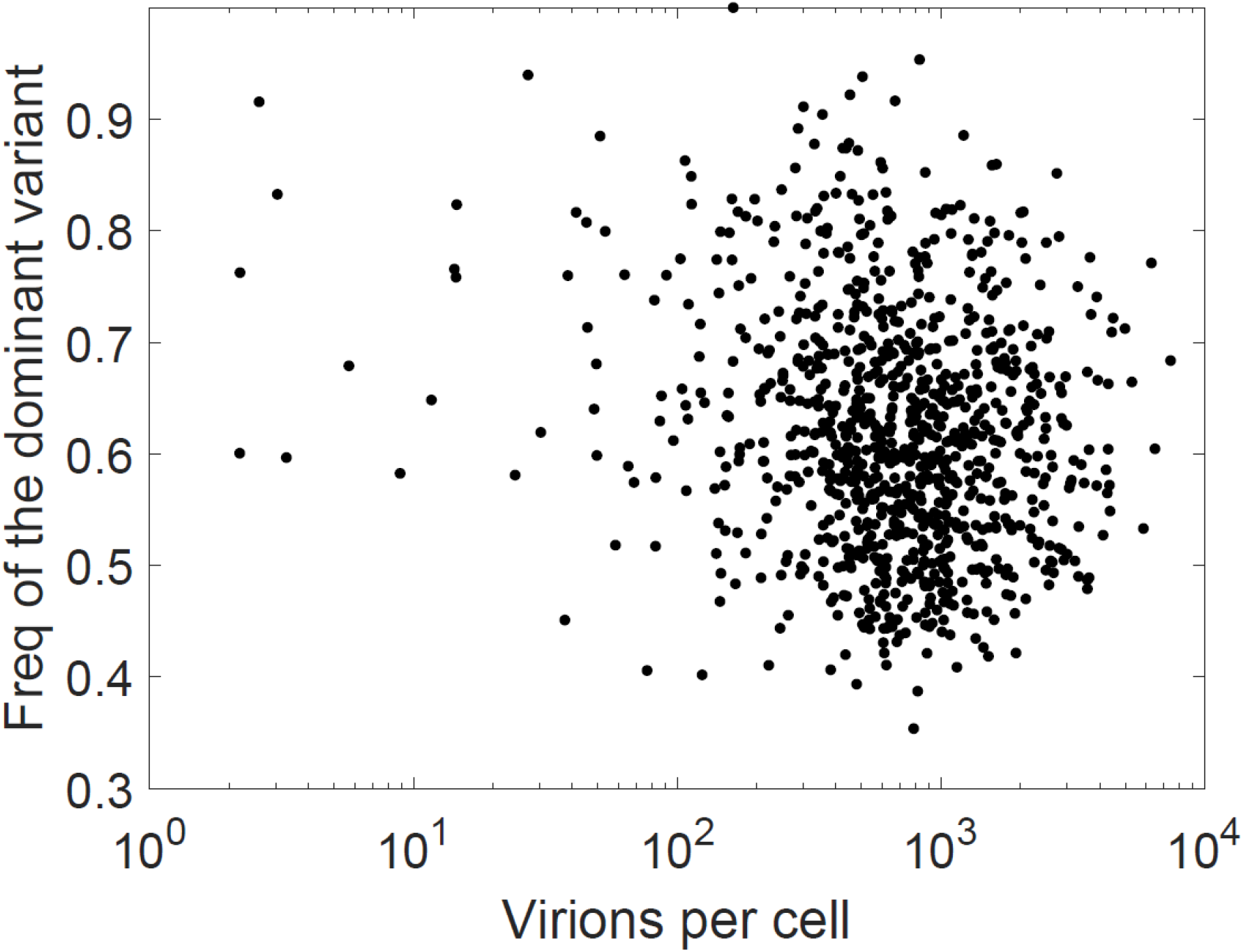
No correlation is observed between the productivity of the infected cells (total virions produced per cell) and the frequency of the most amplified genome of the initial virions (dominant variant), for MOI 10 infections. The pearson correlation coefficient is 0.0185.

**Table S1.** Parameters of the model from (Heldt et al., 2015).

| Parameter | Value | Unit | Description |
| --- | --- | --- | --- |
| kEn | 0.08 | min<sup>-1</sup> | Endocytosis rate. |
| kFus | 0.0535 | min<sup>-1</sup> | Rate of fusion to the endosome membrane. |
| frFus | 0.51 | dimensionless | Fraction of virions that successfully get fused and make it out of the endosome. |
| kImp | 0.1 | min<sup>-1</sup> | Rate of importation to the nucleus. |
| kM | 4166.666667 | nucleotides/min | mRNA transcription rate. |
| K<sub>R</sub> | 1.1E7 | molecules | Polymerase molecules at which transcription rate is half inhibited. |
| kP | 0.023 | min<sup>-1</sup> | Antigenomic synthesis rate. |
| kPr | 1080 | nucleotides/min | Translation rate. |
| 1/Drib | 0.00625 | nucleotides<sup>-1</sup> | Space between two adjacent ribosomes in a single mRNA. |
| kN | 0.231 | min<sup>-1</sup> | Genomic synthesis rate. |
| kBM1 | 2.32E-08 | molecules/min | M1 protein binding rate to RNP. |
| kRel | 6.17E-05 | virions/(molecules x min) | Virion release rate. |
| 1/L1 | 0.000431034 | nucleotides<sup>-1</sup> | Length of segment 1. |
| 1/L2 | 0.000431034 | nucleotides<sup>-1</sup> | Length of segment 2. |
| 1/L3 | 0.000452284 | nucleotides<sup>-1</sup> | Length of segment 3. |
| 1/L4 | 0.000569152 | nucleotides<sup>-1</sup> | Length of segment 4. |
| 1/L5 | 0.000649351 | nucleotides<sup>-1</sup> | Length of segment 5. |
| 1/L6 | 0.000718391 | nucleotides<sup>-1</sup> | Length of segment 6. |
| 1/L7 | 0.000995025 | nucleotides<sup>-1</sup> | Length of segment 7. |
| 1/L8 | 0.001152074 | nucleotides<sup>-1</sup> | Length of segment 8. |
| 1-F_Spl7 | 0.98 | dimensionless | Fraction of spliced mRNA 7 for M1 protein. |
| F_Spl7 | 0.02 | dimensionless | Fraction of spliced mRNA 7 for M2 protein. |
| F_Spl8 | 0.125 | dimensionless | Fraction of spliced mRNA 8 for NEP protein. |
| K_V | 10 | virions | Half maximum proteins amount for virion release rate. |
| kBRdRp | 0.01666667 | molecules/min | Polymerase RNA binding rate. |
| kBNP | 5.0167E-06 | molecules/min | RNA-pol-complex NP binding rate. |
| kExp | 1.6667E-08 | molecules/min | RNP-M1 complex exportation rate to the cytoplasm through the binding of NEP. |
| RNA_deg | 0.606 | min<sup>-1</sup> | Naked RNA degradation rate. |
| RNAPol_deg | 0.07083333 | min<sup>-1</sup> | RNA-pol complex degradation rate. |
| RNPM1c_deg | 0.0015 | min<sup>-1</sup> | RNP-M1 complex in the cytoplasm degradation rate. |
| RNP_deg | 0.0015 | min<sup>-1</sup> | RNP degradation rate. |
| mRNA_deg | 0.0055 | min<sup>-1</sup> | mRNA degradation rate. |
| RNPM1n_deg | 0.0015 | min<sup>-1</sup> | RNP-M1 complex in the nucleus degradation rate. |
| P_deg | 0.000166667 | min<sup>-1</sup> | Protein degradation rate. |
| RdRp | 37 | molecules | Number of polymerases in a virion. |
| HA | 500 | molecules | Number of HA proteins in a virion. |
| NP | 434 | molecules | Number of NP proteins in a virion. |
| NA | 100 | molecules | Number of NA proteins in a virion. |
| M1 | 2932 | molecules | Number of M1 proteins in a virion. |
| M2 | 40 | molecules | Number of M2 proteins in a virion. |
| NEP | 157 | molecules | Number of NEP proteins in a virion. |

## Bibliography

Aggarwal, S., Bradel-Tretheway, B., Takimoto, T., Dewhurst, S., Kim, B., 2010. Biochemical Characterization of Enzyme Fidelity of Influenza A Virus RNA Polymerase Complex. PLoS ONE 5, e10372. 10.1371/journal.pone.0010372

Amato, K.A., Haddock, L.A., Braun, K.M., Meliopoulos, V., Livingston, B., Honce, R., Schaack, G.A., Boehm, E., Higgins, C.A., Barry, G.L., Koelle, K., Schultz-Cherry, S., Friedrich, T.C., Mehle, A., 2022. Influenza A virus undergoes compartmentalized replication in vivo dominated by stochastic bottlenecks. Nat. Commun. 13, 3416. 10.1038/s41467-022-31147-0

Baccam, P., Beauchemin, C., Macken, C.A., Hayden, F.G., Perelson, A.S., 2006. Kinetics of influenza A virus infection in humans. J. Virol. 80, 7590–7599. 10.1128/JVI.01623-05

Brooke, C.B., 2017. Population Diversity and Collective Interactions during Influenza Virus Infection. J. Virol. 91, e01164–17. 10.1128/JVI.01164-17

Brooke, C.B., Ince, W.L., Wrammert, J., Ahmed, R., Wilson, P.C., Bennink, J.R., Yewdell, J.W., 2013. Most influenza a virions fail to express at least one essential viral protein. J. Virol. 87, 3155–3162. 10.1128/JVI.02284-12

Cao, Y., Gillespie, D.T., Petzold, L.R., 2007. Adaptive explicit-implicit tau-leaping method with automatic tau selection. J. Chem. Phys. 126, 224101. 10.1063/1.2745299

Charlesworth, B., 2009. Fundamental concepts in genetics: effective population size and patterns of molecular evolution and variation. Nat. Rev. Genet. 10, 195–205. 10.1038/nrg2526

Cheung, P.P.H., Watson, S.J., Choy, K.-T., Fun Sia, S., Wong, D.D.Y., Poon, L.L.M., Kellam, P., Guan, Y., Malik Peiris, J.S., Yen, H.-L., 2014. Generation and characterization of influenza A viruses with altered polymerase fidelity. Nat. Commun. 5, 4794. 10.1038/ncomms5794

Dolan, P.T., Whitfield, Z.J., Andino, R., 2018. Mapping the Evolutionary Potential of RNA Viruses. Cell Host Microbe 23, 435–446. 10.1016/j.chom.2018.03.012

Ferreri, L.M., Seibert, B., Caceres, C.J., Patatanian, K., Holmes, K.E., Gay, L.C., Cargnin Faccin, F., Cardenas, M., Carnaccini, S., Shetty, N., Rajao, D., Koelle, K., Marr, L.C., Perez, D.R., Lowen, A.C., 2025. Dispersal of influenza virus populations within the respiratory tract shapes their evolutionary potential. Proc. Natl. Acad. Sci. 122, e2419985122. 10.1073/pnas.2419985122

Fitzmeyer, E.A., Gallichotte, E.N., Weger-Lucarelli, J., Kapuscinski, M.L., Abdo, Z., Pyron, K., Young, M.C., Ebel, G.D., 2023. Loss of West Nile virus genetic diversity during mosquito infection due to species-dependent population bottlenecks. iScience 26, 107711. 10.1016/j.isci.2023.107711

Ganti, K., Bagga, A., Carnaccini, S., Ferreri, L.M., Geiger, G., Joaquin Caceres, C., Seibert, B., Li, Yonghai, Wang, L., Kwon, T., Li, Yuhao, Morozov, I., Ma, W., Richt, J.A., Perez, D.R., Koelle, K., Lowen, A.C., 2022. Influenza A virus reassortment in mammals gives rise to genetically distinct within-host subpopulations. Nat. Commun. 13, 6846. 10.1038/s41467-022-34611-z

Ganti, K., Bagga, A., DaSilva, J., Shepard, S.S., Barnes, J.R., Shriner, S., Koelle, K., Lowen, A.C., 2021. Avian Influenza A Viruses Reassort and Diversify Differently in Mallards and Mammals. Viruses 13, 509. 10.3390/v13030509

Garten, R.J., Davis, C.T., Russell, C.A., Shu, B., Lindstrom, S., Balish, A., Sessions, W.M., Xu, X., Skepner, E., Deyde, V., Okomo-Adhiambo, M., Gubareva, L., Barnes, J., Smith, C.B., Emery, S.L., Hillman, M.J., Rivailler, P., Smagala, J., de Graaf, M., Burke, D.F., Fouchier, R.A.M., Pappas, C., Alpuche-Aranda, C.M., López-Gatell, H., Olivera, H., López, I., Myers, C.A., Faix, D., Blair, P.J., Yu, C., Keene, K.M., Dotson, P.D., Boxrud, D., Sambol, A.R., Abid, S.H., St George, K., Bannerman, T., Moore, A.L., Stringer, D.J., Blevins, P., Demmler-Harrison, G.J., Ginsberg, M., Kriner, P., Waterman, S., Smole, S., Guevara, H.F., Belongia, E.A., Clark, P.A., Beatrice, S.T., Donis, R., Katz, J., Finelli, L., Bridges, C.B., Shaw, M., Jernigan, D.B., Uyeki, T.M., Smith, D.J., Klimov, A.I., Cox, N.J., 2009. Antigenic and genetic characteristics of swine-origin 2009 A(H1N1) influenza viruses circulating in humans. Science 325, 197–201. 10.1126/science.1176225

Gerlach, P., Garland, W., Lingaraju, M., Salerno-Kochan, A., Bonneau, F., Basquin, J., Jensen, T.H., Conti, E., 2022. Structure and regulation of the nuclear exosome targeting complex guides RNA substrates to the exosome. Mol. Cell 82, 2505-2518.e7. 10.1016/j.molcel.2022.04.011

Gillespie, D.T., 2001. Approximate accelerated stochastic simulation of chemically reacting systems. J. Chem. Phys. 115, 1716–1733. 10.1063/1.1378322

Goldstein, E., Greene, S.K., Olson, D.R., Hanage, W.P., Lipsitch, M., 2015. Estimating the hospitalization burden associated with influenza and respiratory syncytial virus in New York City, 2003-2011. Influenza Other Respir. Viruses 9, 225–233. 10.1111/irv.12325

Gong, X., Hu, M., Chen, W., Yang, H., Wang, B., Yue, J., Jin, Y., Liang, L., Ren, H., 2021. Reassortment Network of Influenza A Virus. Front. Microbiol. 12. 10.3389/fmicb.2021.793500

Gutiérrez, S., Michalakis, Y., Blanc, S., 2012. Virus population bottlenecks during within-host progression and host-to-host transmission. Curr. Opin. Virol. 2, 546–555. 10.1016/j.coviro.2012.08.001

Heldt, F.S., Frensing, T., Reichl, U., 2012. Modeling the intracellular dynamics of influenza virus replication to understand the control of viral RNA synthesis. J. Virol. 86, 7806–7817. 10.1128/JVI.00080-12

Heldt, F.S., Kupke, S.Y., Dorl, S., Reichl, U., Frensing, T., 2015. Single-cell analysis and stochastic modelling unveil large cell-to-cell variability in influenza A virus infection. Nat. Commun. 6, 8938. 10.1038/ncomms9938

Jacobs, N.T., Onuoha, N.O., Antia, A., Steel, J., Antia, R., Lowen, A.C., 2019. Incomplete influenza A virus genomes occur frequently but are readily complemented during localized viral spread. Nat. Commun. 10, 3526. 10.1038/s41467-019-11428-x

Lee, N., Le Sage, V., Nanni, A.V., Snyder, D.J., Cooper, V.S., Lakdawala, S.S., 2017. Genome-wide analysis of influenza viral RNA and nucleoprotein association. Nucleic Acids Res. 45, 8968–8977. 10.1093/nar/gkx584

Liu, T., Wang, Y., Tan, T.J.C., Wu, N.C., Brooke, C.B., 2022. The evolutionary potential of influenza A virus hemagglutinin is highly constrained by epistatic interactions with neuraminidase. Cell Host Microbe 30, 1363-1369.e4. 10.1016/j.chom.2022.09.003

Lubas, M., Christensen, M.S., Kristiansen, M.S., Domanski, M., Falkenby, L.G., Lykke-Andersen, S., Andersen, J.S., Dziembowski, A., Jensen, T.H., 2011. Interaction profiling identifies the human nuclear exosome targeting complex. Mol. Cell 43, 624–637. 10.1016/j.molcel.2011.06.028

Lumby, C.K., Zhao, L., Breuer, J., Illingworth, C.J., 2020. A large effective population size for established within-host influenza virus infection. eLife 9, e56915. 10.7554/eLife.56915

Martin, B.E., Harris, J.D., Sun, J., Koelle, K., Brooke, C.B., 2020. Cellular co-infection can modulate the efficiency of influenza A virus production and shape the interferon response. PLOS Pathog. 16, e1008974. 10.1371/journal.ppat.1008974

Martin, M.A., Berg, N., Koelle, K., 2024. Influenza A genomic diversity during human infections underscores the strength of genetic drift and the existence of tight transmission bottlenecks. Virus Evol. 10, veae042. 10.1093/ve/veae042

Martínez, F., Sardanyés, J., Elena, S.F., Daròs, J.-A., 2011. Dynamics of a Plant RNA Virus Intracellular Accumulation: Stamping Machine vs. Geometric Replication. Genetics 188, 637–646. 10.1534/genetics.111.129114

Martínez-Alonso, M., Hengrung, N., Fodor, E., 2016. RNA-Free and Ribonucleoprotein-Associated Influenza Virus Polymerases Directly Bind the Serine-5-Phosphorylated Carboxyl-Terminal Domain of Host RNA Polymerase II. J. Virol. 90, 6014–6021. 10.1128/jvi.00494-16

McCrone, J.T., Lauring, A.S., 2018. Genetic bottlenecks in intraspecies virus transmission. Curr. Opin. Virol. 28, 20–25. 10.1016/j.coviro.2017.10.008

Meijers, M., Ruchnewitz, D., Eberhardt, J., Karmakar, M., Luksza, M., Lässig, M., 2024. Concepts and methods for predicting viral evolution. ArXiv 2403.12684v3.

Miller, M., Alvizo, O., Baskerville, S., Chintala, A., Chng, C., Dassie, J., Dorigatti, J., Huisman, G., Jenne, S., Kadam, S., Leatherbury, N., Lutz, S., Mayo, M., Mukherjee, A., Sero, A., Sundseth, S., Penfield, J., Riggins, J., Zhang, X., 2024. An engineered T7 RNA polymerase for efficient co-transcriptional capping with reduced dsRNA byproducts in mRNA synthesis. 10.1039/D4FD00023D

Muñoz-Moreno, R., Martínez-Romero, C., Blanco-Melo, D., Forst, C.V., Nachbagauer, R., Benitez, A.A., Mena, I., Aslam, S., Balasubramaniam, V., Lee, I., Panis, M., Ayllón, J., Sachs, D., Park, M.-S., Krammer, F., tenOever, B.R., García-Sastre, A., 2019. Viral Fitness Landscapes in Diverse Host Species Reveal Multiple Evolutionary Lines for the NS1 Gene of Influenza A Viruses. Cell Rep. 29, 3997-4009.e5. 10.1016/j.celrep.2019.11.070

Nakatsu, S., Sagara, H., Sakai-Tagawa, Y., Sugaya, N., Noda, T., Kawaoka, Y., 2016. Complete and Incomplete Genome Packaging of Influenza A and B Viruses. mBio 7, e01248–16. 10.1128/mBio.01248-16

Nobusawa, E., Sato, K., 2006. Comparison of the mutation rates of human influenza A and B viruses. J. Virol. 80, 3675–3678. 10.1128/JVI.80.7.3675-3678.2006

Parvin, J.D., Moscona, A., Pan, W.T., Leider, J.M., Palese, P., 1986. Measurement of the mutation rates of animal viruses: influenza A virus and poliovirus type 1. J. Virol. 59, 377–383. 10.1128/JVI.59.2.377-383.1986

Pauly, M.D., Procario, M.C., Lauring, A.S., 2017. A novel twelve class fluctuation test reveals higher than expected mutation rates for influenza A viruses. eLife 6, e26437. 10.7554/eLife.26437

Postnikova, Y., Treshchalina, A., Boravleva, E., Gambaryan, A., Ishmukhametov, A., Matrosovich, M., Fouchier, R.A.M., Sadykova, G., Prilipov, A., Lomakina, N., 2021. Diversity and Reassortment Rate of Influenza A Viruses in Wild Ducks and Gulls. Viruses 13, 1010. 10.3390/v13061010

Qu, F., Zheng, L., Zhang, S., Sun, R., Slot, J., Miyashita, S., 2020. Bottleneck, Isolate, Amplify, Select (BIAS) as a mechanistic framework for intracellular population dynamics of positive-sense RNA viruses. Virus Evol. 6, veaa086. 10.1093/ve/veaa086

Rodriguez, A., Pérez-González, A., Nieto, A., 2007. Influenza Virus Infection Causes Specific Degradation of the Largest Subunit of Cellular RNA Polymerase II. J. Virol. 81, 5315–5324. 10.1128/jvi.02129-06

Rüdiger, D., Kupke, S.Y., Laske, T., Zmora, P., Reichl, U., 2019. Multiscale modeling of influenza A virus replication in cell cultures predicts infection dynamics for highly different infection conditions. PLOS Comput. Biol. 15, e1006819. 10.1371/journal.pcbi.1006819

Sardanyés, J., Martínez, F., Daròs, J.-A., Elena, S.F., 2012. Dynamics of alternative modes of RNA replication for positive-sense RNA viruses. J. R. Soc. Interface 9, 768–776. 10.1098/rsif.2011.0471

Sardanyés, J., Solé, R.V., Elena, S.F., 2009. Replication mode and landscape topology differentially affect RNA virus mutational load and robustness. J. Virol. 83, 12579–12589. 10.1128/JVI.00767-09

Schulte, M.B., Draghi, J.A., Plotkin, J.B., Andino, R., 2015. Experimentally guided models reveal replication principles that shape the mutation distribution of RNA viruses. eLife 4, e03753. 10.7554/eLife.03753

Shao, W., Li, X., Goraya, M.U., Wang, S., Chen, J.-L., 2017. Evolution of Influenza A Virus by Mutation and Re-Assortment. Int. J. Mol. Sci. 18, 1650. 10.3390/ijms18081650

Shi, T., Harris, J.D., Martin, M.A., Koelle, K., 2023. Transmission bottleneck size estimation from de novo viral genetic variation. bioRxiv 2023.08.14.553219. 10.1101/2023.08.14.553219

Shi, Y.T., Harris, J.D., Martin, M.A., Koelle, K., 2024. Transmission Bottleneck Size Estimation from De Novo Viral Genetic Variation. Mol. Biol. Evol. 41, msad286. 10.1093/molbev/msad286

Sobel Leonard, A., Weissman, D.B., Greenbaum, B., Ghedin, E., Koelle, K., 2017. Transmission Bottleneck Size Estimation from Pathogen Deep-Sequencing Data, with an Application to Human Influenza A Virus. J. Virol. 91, e00171–17. 10.1128/JVI.00171-17

Suárez, P., Valcárcel, J., Ortín, J., 1992. Heterogeneity of the mutation rates of influenza A viruses: isolation of mutator mutants. J. Virol. 66, 2491–2494. 10.1128/jvi.66.4.2491-2494.1992

Taubenberger, J.K., Kash, J.C., 2010. Influenza Virus Evolution, Host Adaptation and Pandemic Formation. Cell Host Microbe 7, 440–451. 10.1016/j.chom.2010.05.009

Taylor, K.Y., Agu, I., José, I., Mäntynen, S., Campbell, A.J., Mattson, C., Chou, T.-W., Zhou, B., Gresham, D., Ghedin, E., Díaz Muñoz, S.L., 2023. Influenza A virus reassortment is strain dependent. PLoS Pathog. 19, e1011155. 10.1371/journal.ppat.1011155

Teo, Q.W., Wang, Y., Lv, H., Oade, M.S., Mao, K.J., Tan, T.J.C., Huan, Y.W., Rivera-Cardona, J., Shao, E.K., Choi, D., Wang, C., Dargani, Z.T., Brooke, C.B., Velthuis, A.J.W.te, Wu, N.C., 2025. Probing the functional constraints of influenza A virus NEP by deep mutational scanning. Cell Rep. 44. 10.1016/j.celrep.2024.115196

Turrell, L., Lyall, J.W., Tiley, L.S., Fodor, E., Vreede, F.T., 2013. The role and assembly mechanism of nucleoprotein in influenza A virus ribonucleoprotein complexes. Nat. Commun. 4, 1591. 10.1038/ncomms2589

Varble, A., Albrecht, R.A., Backes, S., Crumiller, M., Bouvier, N.M., Sachs, D., García-Sastre, A., tenOever, B.R., 2014. Influenza A Virus Transmission Bottlenecks Are Defined by Infection Route and Recipient Host. Cell Host Microbe 16, 691–700. 10.1016/j.chom.2014.09.020

Wang, C., Forst, C.V., Chou, T.-W., Geber, A., Wang, M., Hamou, W., Smith, M., Sebra, R., Zhang, B., Zhou, B., Ghedin, E., 2020. Cell-to-Cell Variation in Defective Virus Expression and Effects on Host Responses during Influenza Virus Infection. mBio 11, e02880–19. 10.1128/mBio.02880-19

Wang, S., Zhang, T., Hu, M., Tang, K., Sheng, L., Hong, M., Chen, D., Chen, L., Shi, Y., Feng, J., Qian, J., Sun, L., Ding, K., Sun, R., Du, Y., 2023. Deep mutational scanning of influenza A virus neuraminidase facilitates the identification of drug resistance mutations in vivo. mSystems 8, e00670–23. 10.1128/msystems.00670-23

Weaver, S.C., Forrester, N.L., Liu, J., Vasilakis, N., 2021. Population bottlenecks and founder effects: implications for mosquito-borne arboviral emergence. Nat. Rev. Microbiol. 19, 184–195. 10.1038/s41579-020-00482-8

Zwart, M.P., Elena, S.F., 2015. Matters of Size: Genetic Bottlenecks in Virus Infection and Their Potential Impact on Evolution. Annu. Rev. Virol. 2, 161–179. 10.1146/annurev-virology-100114-055135

